# Incorporation of 5 methylcytidine alleviates innate immune response to self-amplifying RNA vaccine

**DOI:** 10.1101/2023.11.01.565056

**Authors:** Mai Komori, Amber L. Morey, Andrés A. Quiñones-Molina, Josiane Fofana, Luis Romero, Elizabeth Peters, Kenta Matsuda, Suryaram Gummuluru, Jonathan F. Smith, Wataru Akahata, Hisashi Akiyama

## Abstract

In order to improve vaccine effectiveness and safety profile of existing synthetic RNA-based vaccines, we have developed a self-amplifying RNA (saRNA)-based vaccine expressing membrane-anchored receptor binding domain (RBD) of SARS-CoV-2 S protein (S-RBD) and have demonstrated that a minimal dose of this saRNA vaccine elicits robust immune responses. Results from a recent clinical trial with 5-methylcytidine (5mC) incorporating saRNA vaccine demonstrated reduced vaccine-induced adverse effects while maintaining robust humoral responses. In this study, we investigate the mechanisms accounting for induction of efficient innate and adaptive immune responses and attenuated adverse effects induced by the 5mC-incorporated saRNA. We show that the 5mC-incorporating saRNA platform leads to prolonged and robust expression of antigen, while induction of type-I interferon (IFN-I), a key driver of reactogenicity, is attenuated in peripheral blood mononuclear cells (PBMCs), but not in macrophages and dendritic cells. Interestingly, we find that the major cellular source of IFN-I production in PBMCs is plasmacytoid dendritic cells (pDCs), which is attenuated upon 5mC incorporation in saRNA. In addition, we demonstrate that monocytes also play an important role in amplifying proinflammatory responses. Furthermore, we show that the detection of saRNA is mediated by a host cytosolic RNA sensor, RIG-I. Importantly, 5mC-incorporating saRNA vaccine candidate produced robust IgG responses against S-RBD upon injection in mice, thus providing strong support for the potential clinical use of 5mC-incorporating saRNA vaccines.

## Introduction

Despite the success of mRNA-based vaccines to combat the SARS-CoV-2 pandemic, there remains a need to improve both vaccine effectiveness and safety profile. One of the major challenges for the current mRNA vaccine formulations is vaccine reactogenicity. Many vaccinees have experienced mild to severe systemic adverse effects such as fever, chills and joint pains (reviewed in ^1,2^). Although the precise mechanisms that underlie severe systemic adverse effects are not yet known, it is assumed that excessive innate immune reactions to components of vaccines causes these adverse effects ^3^. Therefore, it is important to understand how synthetic RNA-based vaccines induce excessive innate immune responses to enable the development of strategies to reduce reactogenicity. It is well known that synthetic exogenous RNA triggers robust innate immune responses via pathogen recognition receptors such as TLRs ^4^, which had previously prevented the development of synthetic mRNA as a vaccine platform. Unlike synthetic RNA, endogenous RNAs found in cells are post-transcriptionally modified at many residues ^5-7^, and this posttranscriptional RNA modifications were postulated to be one of the ways that host machinery distinguishes self- and non-self RNAs. In 2005, Kariko and colleagues reported that the replacement of uridines with pseudouridine (Ψ) in synthetic RNAs made them non-immunostimulatory ^8^, and these seminal findings paved the way to the use of synthetic mRNAs as a vaccine platform in vivo.

Antigen production and immunogenicity are generally dependent on the dose of mRNAs delivered. Although mRNA vaccines have been optimized to increase stability and antigen expression, mRNA still undergoes degradation ^9^. Therefore, to elicit efficient and prolonged immune responses, COVID19 vaccines require multiple injections and high doses mRNA (30-100 µg mRNA per dose). To develop a more efficient RNA vaccine, we have utilized self-amplifying RNA (saRNA)-based vaccine candidates expressing membrane-anchored receptor binding domain (RBD) of SARS-CoV-2 S protein (S-RBD)^10^. This single-cycle vector system utilizes an alphavirus RNA amplification system derived from the Venezuelan Equine Encephalitis Virus (VEEV)-based replicon expression vector ^11,12^. VEEV has a single-stranded positive-sense RNA as its genome, and the genome contains two open reading frames (ORFs). The 5’-proximal ORF encodes the alphavirus nonstructural proteins (nsPs)1-4 which provide the RNA-dependent RNA polymerase (RdRp) and transcriptase functions. ^16-23^. The 3’ proximal ORF is transcribed from a subgenomic promoter and encodes the viral structural proteins. For the saRNA vectors, the ORF for the structural proteins was replaced with a sequence encoding a membrane-anchored S-RBD. This vector expresses the VEEV replication machinery to replicate and transcribe the saRNA, resulting in efficient expression of the gene(s) of interest. Due to this self-amplification process, the level and duration of expression of target antigens are expected to be higher and more durable than that observed with nonreplicating mRNA vaccine platforms ^10^. One clinical trial using an saRNA vector expressing the whole SARS S protein demonstrated that 5 µg of the saRNA induced significant amounts of neutralizing antibodies ^13^. In addition, in a Phase I study, 3 µg of saRNA encoding membrane anchored S-RBD as the antigen elicited robust IgG responses which were comparable to those induced by 30 µg BNT162b2 vaccine ^14^. Importantly, a separate phase 1 study using saRNA encoding a membrane-anchored S-RBD containing 5mC showed reduced systemic reactogenicity while maintaining IgG responses ^15^. Despite the promising clinical data with saRNA-based vaccines, the mechanisms of how saRNA vaccines induce reactogenicity and how incorporation of modified RNA bases attenuates systemic reactogenicity have remained unclear.

In this study, we sought to determine the mechanisms by which innate immune response, a key driver of reactogenicity, induced by LNP-formulated saRNA vaccine candidates targeting S-RBD. We found that both mRNA and saRNA induced innate immune responses in cell lines, and there were cell-type and vaccine-format specific responses. In PBMCs, we detected a robust type-I interferon (IFN-I) response to LNP-formulated saRNA. Interestingly, incorporation of 5mC attenuated IFN-I responses in PBMCs, but not in monocyte-derived macrophages and dendritic cells. While the major source of IFN-I production was plasmacytoid dendritic cells (pDC), monocytes also played an important role in amplifying proinflammatory responses. The detection of saRNA was mediated by a host cytosolic RNA sensor, RIG-I. Importantly, a 5mC-incorporated saRNA vaccine candidate produced a robust IgG response against S-RBD upon injection in mice.

## Results

### saRNA provides robust and prolonged expression of membrane anchored S-RBD

For the saRNA vector used in this study, the ORF for structural proteins was replaced with a sequence encoding a membrane-anchored S-RBD (Fig. 1A). To test the impact of RNA base modification on host innate and adaptive immune responses to synthetic RNAs, we synthesized mRNA/saRNA with native RNA bases (native), N1-methyl-pseudouridine (N1mΨ) instead of uridine, or 5-methyl-cytidine (5mC) instead of cytidine. To examine expression of S-RBD from mRNA or saRNA, two indicator cell lines were employed. HEK293-ISRE-luc cells (hereafter 293-ISRE) were engineered to express luciferase reporter gene under the control of an interferon-stimulated response element (ISRE) ^24,25^. The second cell line, CHME 5xISRE Nluc (hereafter CHME-ISRE), was created by transducing a human microglia cell line CHME3 (also known as HMC3 authenticated by ATCC ^26,27^) with a lentivector expressing Nluc under the control of five tandem ISRE repeats. 293-ISRE cells or CHME-ISRE cells were incubated with LNPs containing mRNA (native, N1mΨ, 5mC) or saRNA (native, N1mΨ, 5mC) encoding S-RBD, and S-RBD expression was analyzed by flow cytometry (FCM). FCM revealed prolonged expression of S-RBD from saRNA in both cell types (Fig. 1B). In contrast to mRNA-dependent S-RBD expression in 293-ISRE cells, S-RBD expression from saRNA was maintained until day 4 post addition (native and 5mC), while expression of S-RBD from mRNA was markedly reduced in CHME-ISRE cells by day 4 (Fig. 1B). In CHME-ISRE cells, although expression of S-RBD from saRNA did not change on day 1 and day 2, expression of S-RBD from mRNA was already reduced on day 2 (Fig. 1B). Interestingly, replacement of uridine with N1mΨ led to the loss of S-RBD expression from saRNA in both cell types, while mRNA-N1mΨ was able to express S-RBD (Fig. 1B). These results demonstrate that the saRNA platform provides efficient and prolonged antigen expression, and that 5mC is compatible with the VEEV replication machinery unlike N1mΨ.

**Fig. 1.**
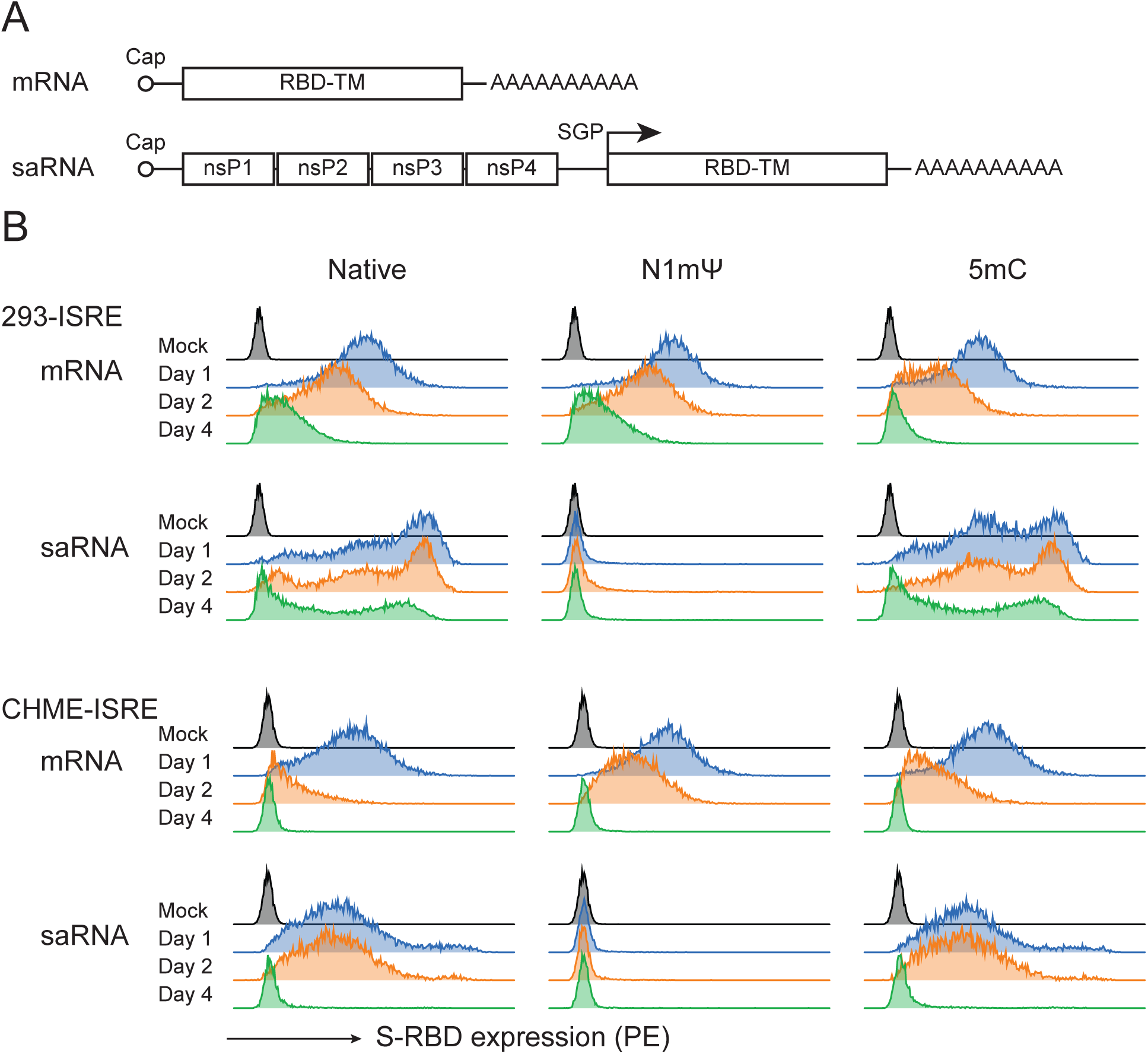
saRNA provides robust and prolonged expression of membrane-anchored S-RBD. (A) Schematics of mRNA vector and saRNA vector expressing membrane anchored SARS-CoV-2 S-RBD (RBD-TM). SGP: subgenomic promoter. (B) S-RBD expression profiles in 293-ISRE cells and CHME-ISRE cells. Cells were incubated with LNPs containing mRNA (native, N1mΨ, 5mC) or saRNA (native, N1mΨ, 5mC) and surface S-RBD expression was analyzed by flow cytometry on day 1, 2 and 4 post incubation. Representative histograms are shown. Mock: cells without LNPs.

### LNP-formulated mRNA and saRNA induce innate immune activation in a cell type specific manner

To investigate innate immune responses induced by LNP-formulated mRNA and saRNA, we utilized three different indicator cell lines: 293-ISRE, CHME-ISRE, and THP-1 Dual cells (NF-κB-SEAP IRF-Luc reporter monocytes, Invivogen). These cell lines were incubated with LNPs containing mRNA or saRNA encoding S-RBD (native, N1mΨ, 5mC), and luciferase/Nluc expression was measured 24-48 hours post stimulation. In 293-ISRE cells, mRNA did not induce any luciferase signals, while saRNA (native and 5mC) induced robust innate immune responses in a dose dependent manner (Fig. 2A and S1A). In CHME-ISRE cells, on the other hand, both mRNA-native and saRNA-native induced innate immune activation (Fig. 2B and S1B). Interestingly, incorporation of m5C attenuated innate immune activation induced by mRNA but this was not observed in saRNA. In THP-1 Dual cells, both mRNA-native and saRNA-native led to induction of IFN responses, however, 5mC incorporation dampened innate activation in both vaccine formats (Fig. 2C). Replacement of uridine with N1mΨ abrogated innate immune activation in all three cell lines with both mRNA and saRNA, in agreement with the previous findings^8^. These results suggest that cell-type specific host mechanisms recognize modified RNA bases in synthetic RNAs and modulate the sensing mechanism of non-host RNAs.

**Fig. 2.**
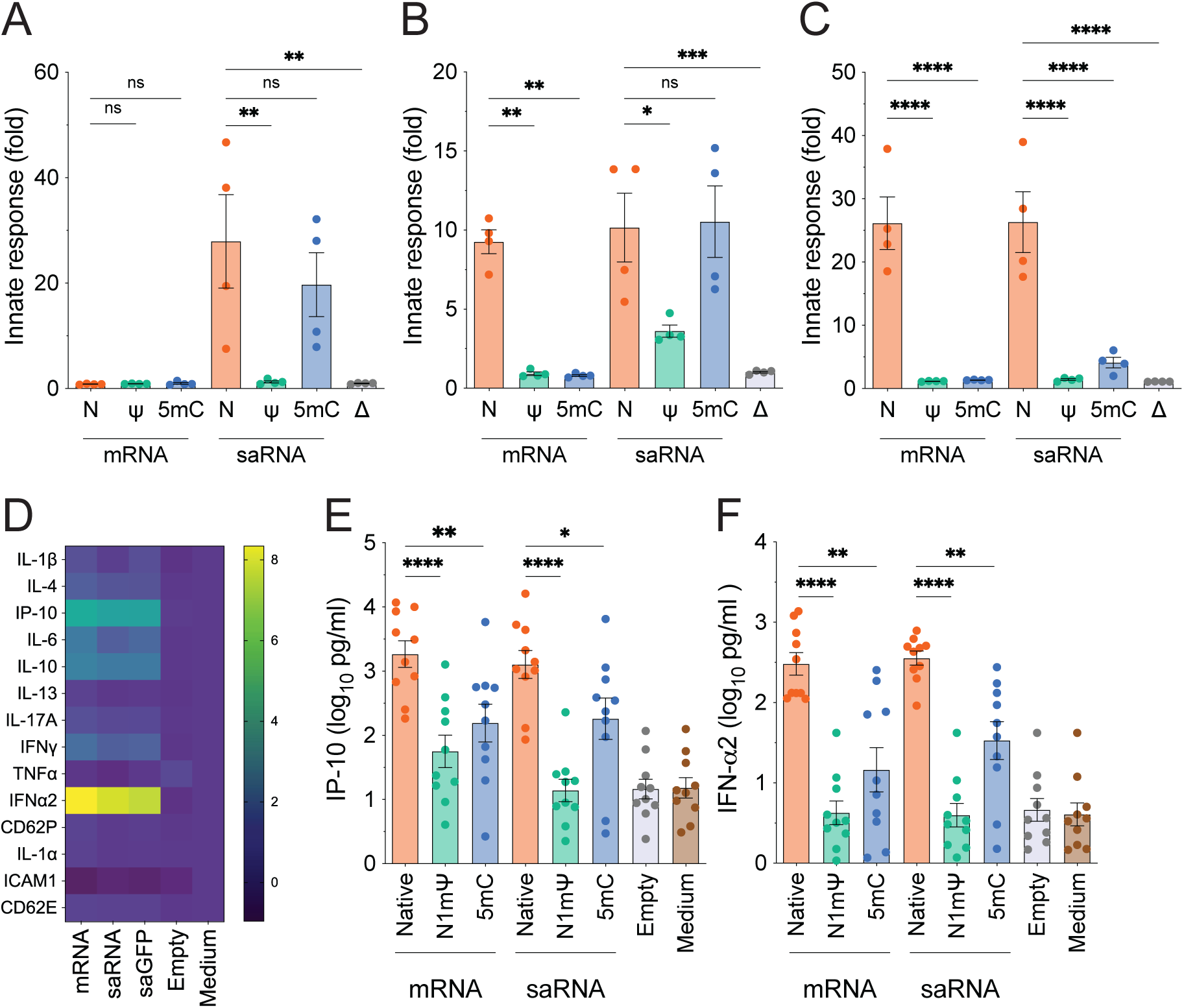
LNP-formulated mRNA and saRNA induce innate immune activation, which is attenuated by the incorporation of modified RNA bases in a cell type specific manner. (A) 293-ISRE cells, (B) CHME-ISRE cells, or (C) THP-1 Dual cells were incubated with LNPs containing mRNA or saRNA with native bases (N), N1mΨ (Ψ), or 5mC, or empty LNPs (Δ) at 50 ng/ml (A and B) or 500 ng/ml (C), and luciferase or Nluc activity was quantified 24-48 hours post stimulation. Values were normalized to that of medium only control. The data represent the means +/- SEM, and each dot represents an individual experiment. (D) PBMCs from healthy donors were incubated with mRNA-native (mRNA), saRNA-native (saRNA), saRNA expressing GFP (saGFP), or empty LNPs (Empty) at 50 ng/ml, culture supernatants were harvested 24 hours post initiation of culture, and cytokines released into the sups were quantitated with multiplex cytokine assay (Luminex). Values were normalized to those of medium only control, and fold change in log_10_ is shown. n=10. (E and F) PBMCs were incubated with mRNA (native, N1mΨ, 5mC), saRNA (native, N1mΨ, 5mC), or empty LNPs 50 ng/ml for 24 hours, and cytokines released into the sups were quantitated with multiplex cytokine assay (LegendPlex). The graph shows the means +/- SEM. Each dot represents an individual experiment with cells from a different donor. One-way ANOVA, *: *p*<0.05, **: *p*<0.01, ***: *p*<0.001, ****: *p*<0.0001, ns: not significant.

### Incorporation of modified RNA bases attenuated innate immune responses to LNP-formulated mRNA and saRNA

To examine innate immune sensing of synthetic RNAs in primary cells, we isolated peripheral blood mononuclear cells (PBMCs) from healthy donors and cultured them with LNPs containing 50 ng/ml mRNA-native, saRNA-native, saRNA-GFP (encoding GFP instead of S-RBD), or empty LNPs for 3 days. Culture supernatants were harvested daily until Day 3, and cytokine release into the supernatants were measured by a multiplex cytokine assay. We observed robust type I interferon (IFN-I) responses (IFN-⍺2) and proinflammatory cytokine IP-10 production in LNP-treated PBMC cultures (Fig 2D). The induction of IFN-I responses was not antigen-dependent since saRNA-native (antigen: S-RBD) and saRNA-GFP (antigen: GFP) both induced similar levels of IFN-⍺2 and IP-10 production. Induction of IFN-a2 decreased over time, while IP-10 secretion increased over time (Fig. S2A and S2B).

Next, we tested whether the incorporation of N1mΨ or 5mC affects innate immune responses in PBMCs. Multiplex cytokine assay revealed that IP-10 and IFN-⍺2 secretion were severely attenuated in PBMCs cultured with mRNA-N1mΨ and saRNA-N1mΨ (Fig. 2E and F). Interestingly, incorporation of 5mC significantly reduced production of IP-10 and IFN-⍺2 in response to both mRNA and saRNA (Fig. 2E and F).

### Role of monocytes, macrophages and dendritic cells in response to LNP-formulated saRNA

To further investigate the mechanism of innate immune activation in primary cells, we tested innate responses to synthetic RNAs in various cell types that potentially take up LNP-saRNA and trigger innate immune responses. For the goal, we updated our saRNA platform to the saRNA-PADRE vector which encodes the pan HLA-DR-binding epitope (PADRE) at the C terminus of S-RBD (Fig. S3A). PADRE encodes a short stretch of peptides that binds to most common leukocyte antigen-DR isotypes, and are designed to enhances adaptive immunity to S-RBD in an MHC-II-restricted manner ^28^. Since this platform is currently in clinical testing ^15^, the saRNA-PADRE vector is clinically more relevant. Note that there is no difference between saRNA and saRNA-PADRE (native and 5mC) in induction of innate immune responses in PBMCs (Fig. S3B and 3C).

Monocytes have been shown to be one of the cell types that efficiently take up RNA vaccines in non-human primates ^29^. To test whether monocytes are responsible for innate immune activation and are differentially responsive to modified RNA bases in synthetic RNAs, CD14^+^ monocytes, isolated from PBMCs, were cultured with synthetic RNAs and stained for CD169 (a myeloid specific surface receptor that is induced by IFN-I and III ^24,30^). We found that CD169 was quickly and robustly up-regulated in CD14^+^ monocytes, while saRNA-5mC delayed monocyte activation and saRNA-N1mΨ did not activate monocytes (Fig. 3A). Next, we tested if monocytes were the producer of IP-10 and IFN-⍺2 in saRNA-exposed PBMCs. CD14^+^ monocytes were isolated from PBMCs and cultured with LNPs containing saRNAs. Interestingly, monocytes did not produce IP-10 or IFN-⍺2 in response to saRNA (Fig. 3B and 3C), suggesting that some cell types other than monocytes in PBMCs induce IFN-I responses. In support of this hypothesis, when PBMCs were cultured with LNP-saRNA-native in the presence of IFN-I decoy receptor B18R ^31^, IP-10 secretion was almost completely inhibited (Fig. 3D), suggesting IFN-I from other cell types in PBMCs activates monocytes. Interestingly, CD14-depleted PBMCs also showed markedly reduced expression of IP-10, suggesting that monocytes amplify inflammatory responses in response to IFN-I in a paracrine manner (Fig. 3E).

**Fig. 3.**
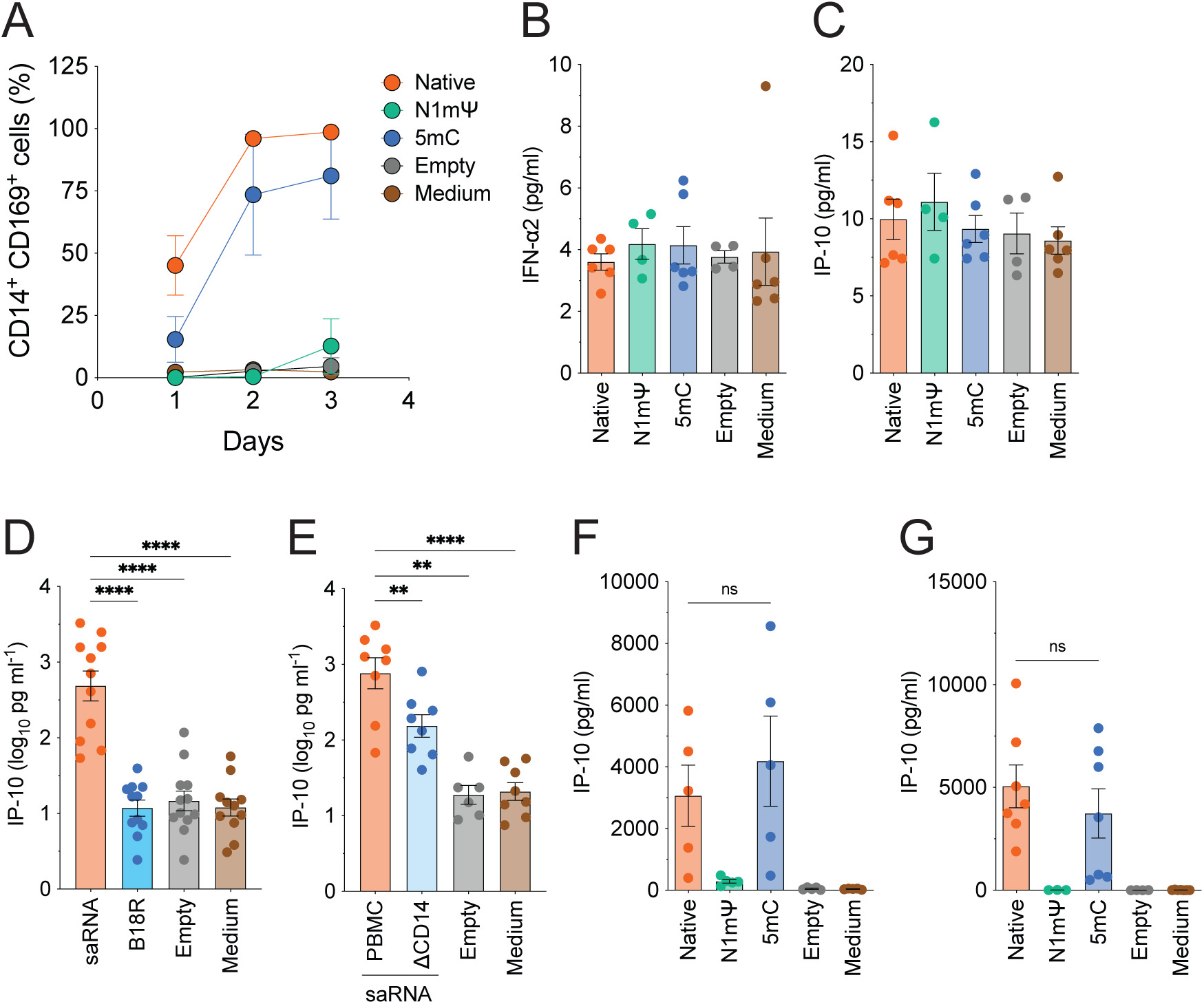
Role of monocytes, macrophages and dendritic cells in response to LNP formulated saRNA. (A) PBMCs were cultured with LNPs containing saRNA-PADRE (native, N1mΨ, 5mC), or empty LNPs (Empty) at 50 ng/ml and stained for live/dead, CD14 and CD169 on day 1, 2, and 3 post initiation of culture and analyzed by flow cytometry. The percentages of live CD169^+^ monocytes are shown. The graph shows the means +/- SEM. n=4. (B and C) Monocytes isolated from PBMCs were cultured with LNPs containing saRNA-PADRE (native, N1mΨ, 5mC), or empty LNPs (Empty) at 50 ng/ml for 1 day, and (B) IFN-⍺2 and (C) IP-10 released into the culture were analyzed by multiplex cytokine assay (LegendPlex). (D) PBMCs were cultured with LNPs containing saRNA-PADRE-native in the presence (1 µg/ml) or absence of B18R, an IFNAR decoy receptor, and IP-10 release on day 1 was measured with LegendPlex. (E) PBMCs or CD14^+^ cell-depleted PBMCs (ΔCD14) were cultured with LNPs containing saRNA-PADRE-native at 50 ng/ml and IP-10 production on day 1 was measured by LegendPlex. (F) MDMs and (G) MDDCs were generated from PBMCs and incubated with LNPs containing saRNA-PADRE (native, N1mΨ, 5mC), or empty LNPs (Empty) at 50 ng/ml and IP-10 release into the culture supernatants on day 3 was measured by ELISA (F) or LegendPlex (G). (B-G) Each dot represents an individual experiment with cells from a different donor. One-way ANOVA, **: *p*<0.01, ****: *p*<0.0001, ns: not significant.

In addition to monocytes, we have tested macrophages and dendritic cells as tissue resident cells which respond to vaccines at the injection site. We isolated monocytes from PBMCs using magnetic beads and differentiated into macrophages (monocyte-derived macrophages, MDMs) or dendritic cells (monocyte-derived dendritic cells, MDDCs) as previously reported ^24^ and used as a surrogate for tissue-resident macrophages and DCs, respectively. MDMs or MDDCs were incubated with LNP-saRNA-native, -N1mΨ or -5mC, and cytokine secretion in the culture sups was measured. In alignment with the data in PBMCs, saRNA-native induced IP-10 in both cell types and N1mΨ incorporation inhibited the secretion (Fig 3F and G). Interestingly, we did not observe significant differences between saRNA-native and saRNA-5mC in IP-10 production in MDMs and MDDCs (Fig. 3F and 3G), suggesting that there is another different cell type other than monocytes, macrophages and DCs which contributes to cell-type specific mechanism by which 5mC attenuates induction of innate immune responses in PBMCs.

### Plasmacytoid dendritic cells (pDCs) induces robust IFN-I responses to saRNA, that is reduced by 5mC incorporation

pDCs are only a minor population in PBMCs, however, they secrete a robust amount of IFN-I upon stimulation with pathogens such as viruses ^32^. It has been shown that upon transfection of synthetic RNAs with 5mC in pDCs, there was less cytokine release ^8^. Therefore, we sought to determine the role of pDCs in sensing RNA vaccine candidates. pDCs were isolated from PBMCs by magnetic beads and incubated with LNPs containing saRNA-native, -N1mΨ, or -5mC, and cytokine secretion was measured by multiplex cytokine assay. Alternatively, pDC were depleted from PBMCs prior to incubation with these LNPs. We found that pDCs secreted robust amounts of IFN-I such as IFN-⍺2 (Fig. 4A and S4A) and IFN-β (Fig. S4B) as well as other proinflammatory cytokines including IFN-λ1 (Fig. S4A and S4C). Moreover, pDC-depleted PBMCs were severely attenuated in IFN-I responses upon stimulation with saRNA (Fig. 4B), suggesting that pDCs are the primary IFN-I secreting cell in PBMCs upon stimulation with synthetic RNAs in PBMCs. Interestingly, the incorporation of 5mC in saRNA attenuated secretion of IFN-I and other cytokines from pDCs (Fig. 4A, S4A-C). It is most likely, therefore, that the difference in sensing of synthetic RNAs with cytidine or 5mC in PBMCs was reflected by the impaired response to 5mC-containing synthetic RNAs in pDCs. In contrast, pDC produced only a small amount of IP-10 upon stimulation with saRNA and there was no difference between saRNA-native and saRNA-5mC in IP-10 production (Fig. 4C), suggesting that pDC hardly contributed to the production of IP-10 in PBMCs upon stimulation with saRNA. Lastly, to test the possibility that monocytes produce IP-10 in response to IFN-I produced from pDCs upon stimulation with synthetic RNAs, we co-cultured monocytes and pDCs in the presence or absence of LNP-formulated saRNA. We found that, although pDCs did not produce a high level of IP-10 compared to the whole PBMCs (Fig. 4C), cytokines, most likely IFN-I, produced from pDCs stimulated monocytes, resulted in a robust secretion of IP-10 (Fig. 4D). In summary, our data demonstrate the importance of pDCs in responding to synthetic RNAs, and the incorporation of 5mC significantly affects this response.

**Fig. 4.**
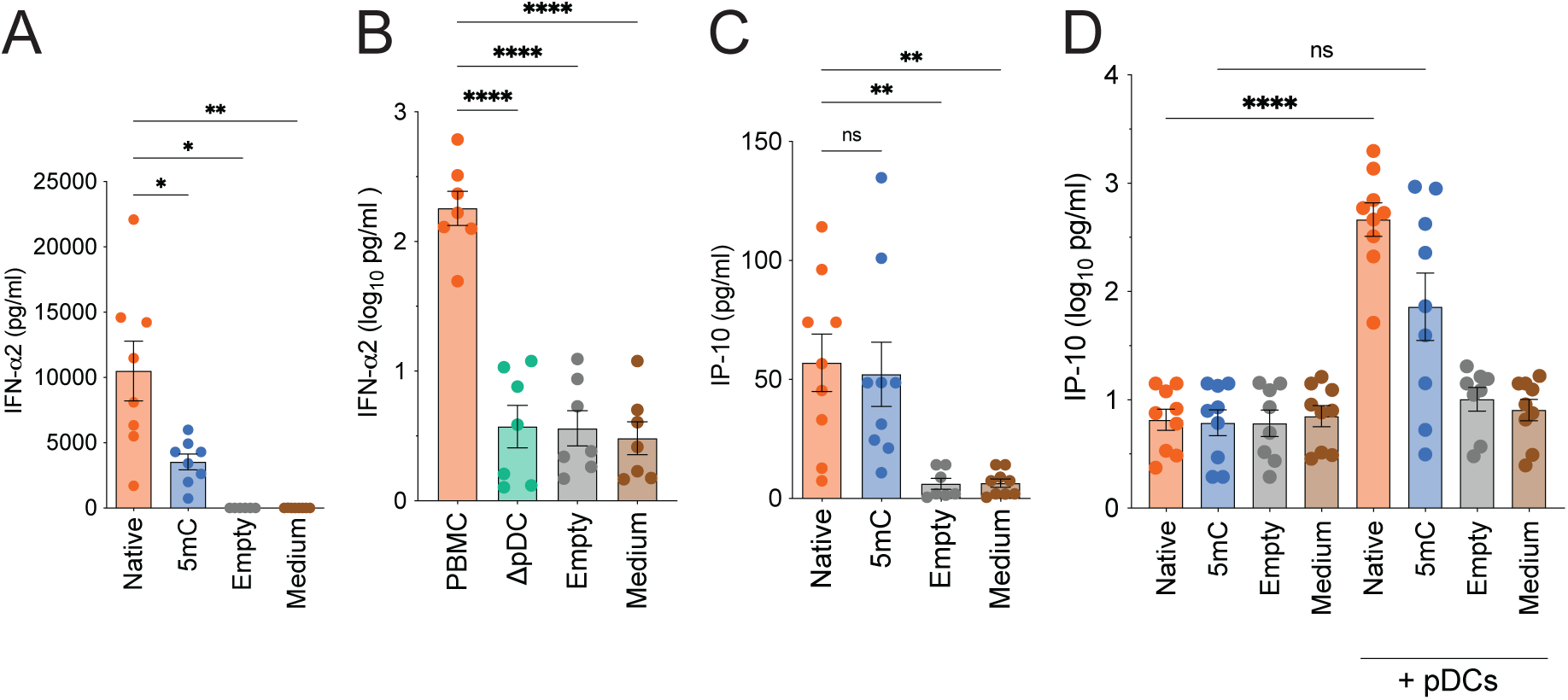
pDCs induce robust IFN-I responses to saRNA, which is reduced by m5C incorporation. (A) pDCs isolated from healthy donor PBMCs were incubated with LNPs containing saRNA-PADRE (native or 5mC), or empty LNPs at 50 ng/ml for 1 day, and the production of IFN-⍺2 was measured with LegendPlex. (B) PBMCs or pDC-depleted PBMCs (ΔpDC) were incubated with LNPs containing saRNA-PADRE-native, or empty LNPs at 50 ng/ml for 1 day, and the production of IFN-⍺2 was measured with LegendPlex. (C) pDCs were incubated with LNPs containing saRNA-PADRE (native or 5mC), or empty LNPs at 50 ng/ml for 1 day, and the production of IP-10 was measured with LegendPlex. (D) Monocytes or monocyte/pDC co-culture was incubated with LNPs containing saRNA-PADRE-native, and IP-10 production was examined with LegendPlex. Each dot represents an individual experiment using cells from a different donor. The graph shows the means +/- SEM. One-way ANOVA, *: *p*<0.05, **: *p*<0.01, ****: *p*<0.0001, ns: not significant.

### Innate immune response to saRNA is mediated by RIG-I

Cells are equipped with many intrinsic mechanisms to detect foreign nucleic acids such from pathogens and amount a defense against these invading pathogens. These include endosomal DNA/RNA sensors such as TLRs and intracellular cytoplasmic nucleic acid sensors such as RIG-I/MDA5 (RNA sensors) and cGAS (DNA sensor). It has been shown that synthetic mRNA is sensed by RIG-I. To investigate the role of cytosolic sensors RIG-I and MDA5 in sensing saRNA, we knocked down expression of these sensors as well as MAVS, which transduces signals from RIG-I and MDA5 to induce IFN-I and pro-inflammatory cytokines ^33^, in 293-ISRE cells and CHME-ISRE cells using lentivectors expressing shRNA against the target mRNAs. We also established knockdown (KD) cell lines of STING, which is a signal transducer downstream of DNA sensing ^34^ Cells expressing shRNA to scrambled sequences served as negative controls. KD efficiency was validated with qRT-PCR (Fig. S5A and S5B) and western blotting analysis (Fig. S5C). STING expression in 293-ISRE cells was below the detection limit of western blotting, however, the expression of other proteins was confirmed and successfully reduced by shRNA (Fig. S5C). These KD cell lines were incubated with increasing doses of LNPs containing saRNA-native, and innate immune activation was measured. In 293-ISRE cells, KD of RIG-I significantly reduced induction of signals and KD of MAVS completely abrogated innate immune activation by both types of synthetic RNAs, demonstrating that the RIG-I-MAVS axis is important for mRNA and saRNA sensing (Fig. 5A). In CHME-ISRE, KD of RIG-I reduced innate immune activation, indicative of an important role of RIG-I in saRNA sensing in this cell type as well (Fig. 5B). KD of MAVS in CHME-ISRE cells did not impact on saRNA-mediated innate immune activation for an unknown reason. These results indicate that RIG-I is the key player in sensing saRNAs.

**Fig. 5.**
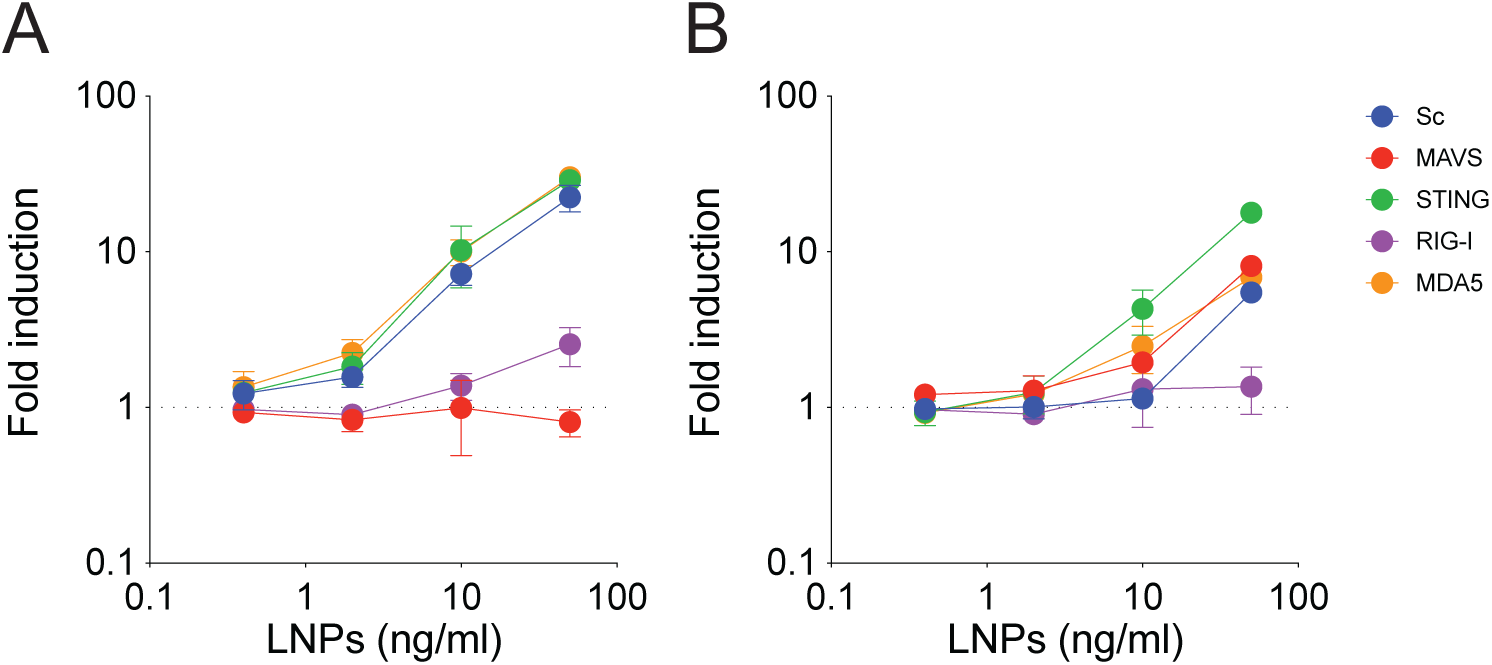
Innate immune response to saRNA is mediated by RIG-I. Innate immune activation in (A) 293-ISRE or (B) CHME-ISRE cells with KD of the indicated protein. Sc: Scrambled shRNA expressing cells. Cells were incubated with increasing amounts of LNPs containing saRNA-PADRE-native and luciferase or NLuc activity was measured on day 1 (D), or day 2 (E) post saRNA addition. Values were normalized to those from media-only control. Representative graphs are shown (n=5).

### 5mC-incorporated saRNA vaccine elicits a robust humoral response in vivo

Immunogenicity of the LNP-saRNA comprising native, N1mΨ, or 5mC bases were assessed in mice. Female Balb/c mice (6 mice per group) were immunized with 10µg of LNP-saRNA at day 0 and 28 and blood collection was performed at day 0, 14, 21, 29, 42, and 56 post immunization (Fig. 6A). ELISA was performed to test circulating IgG antibody titers against SARS-CoV2 spike protein (Gamma). The antibody titers in mice immunized with native and 5mC incorporated LNP-saRNA were comparable at all timepoints up to day 56 (Fig. 6B and C). On the other hand, mice immunized with mN1mΨ incorporated LNP-saRNA showed significantly reduced IgG titers at all time points we have assessed (Fig. 6B and C). These results demonstrate that the incorporation of 5mC in the saRNA platform does not impair its ability to present antigens to the immune system and results in prolonged induction of humoral responses to the antigen, consistent with our *in vitro* results.

**Fig. 6.**
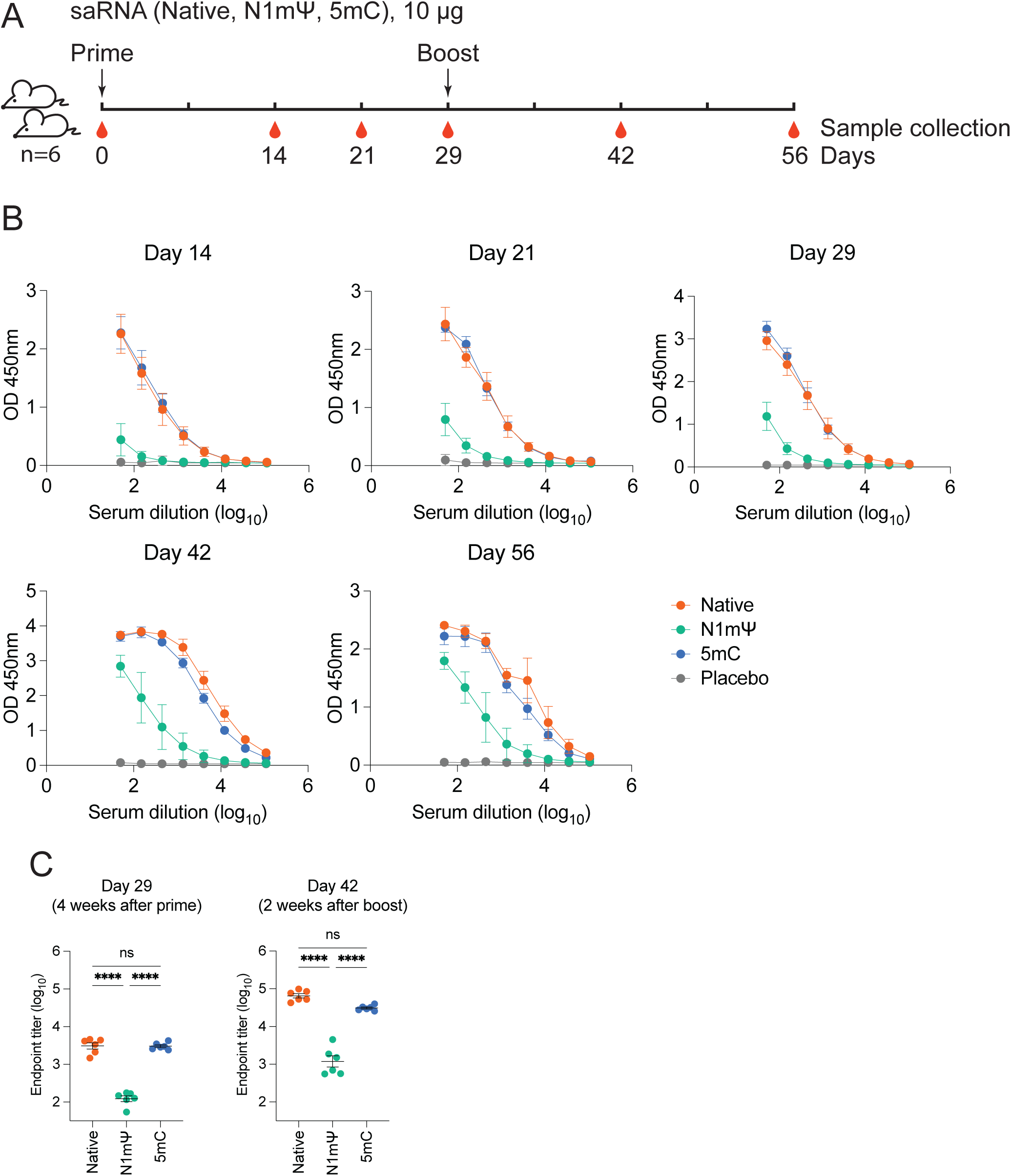
saRNA induces robust humoral responses against S-RBD in mice. (A) Vaccination and sample collection schedule. Female BALB/c mice (n=6 per group) were immunized intramuscularly twice at a four-week interval with 10µg of LNP-saRNA consists of native, N1mΨ, or 5mC. (B) Antibody titers in serum were measured by ELISA. The graph shows the means +/- SD. (C) Plot represents individual endpoint titers with the means +/- SEM. One-way ANOVA, ****: *p*<0.0001, ns: not significant.

## Discussion

The seminal findings by Kariko and colleagues that incorporation of modified RNA bases such as Ψ prevents excessive innate immune responses to synthetic RNA ^8^ and enhanced antigen expression ^35^ has led to the success of mRNA-based vaccines. Similar efforts to enhance immunogenicity and attenuate innate immunity in saRNA by incorporating modified uridines have been unsuccessful ^36,37^. Therefore, it has been thought that alpha virus RdRp is incompatible with modified RNA bases, leading to lack of antigen expression^38^. In this study, we have tested cytosine substitution with 5mC in the saRNA platform and found, unlike uridine substitution, that incorporation of 5mC is compatible with saRNA function and leads to enhanced and prolonged expression of the antigen relative to the conventional mRNA platform. Interestingly, m5C incorporation into saRNA attenuates IFN-I and proinflammatory responses in PBMCs mainly due to the limited activation of the IFN-I pathway in pDCs. Although further reactogenicity studies of saRNA-m5C in animal models are currently in progress, we have already observed attenuated systemic reactions such as fevers in a human clinical trial conducted with saRNA-m5C vaccines ^15^. While we were preparing this manuscript, a preprint by McGee and colleagues has also shown that cytosine modification enables saRNA to show enhanced and prolonged antigen expression and attenuated IFN-I responses ^39^. Findings in this preprint and in our *in vitro* studies, in animals, and in a clinical trial strongly demonstrate that the incorporation of modified nucleoside such as 5mC improves saRNA vectors both with respect to reduced reactogenicity and with respect to enhanced and sustained immunogenicity.

We investigated the mechanisms of innate immune responses to mRNA and saRNA in two cell lines and have shown that RIG-I plays an important role in sensing in both cell types. However, there are also several notable differences between these cell types and between vaccine platforms. In 293-ISRE cells, mRNA did not induce innate activation while saRNA induced a robust signal. saRNA vectors produce double-stranded RNA (dsRNA) during their amplification process (saRNA produces negative-sense RNA as a template for both full-length positive sense viral RNA synthesis and for transcription of subgenomic mRNA). Therefore, it is plausible that RIG-I does not respond to the incoming synthetic RNAs but responds to dsRNA of saRNA in 293-ISRE cells. In contrast, in CHME-ISRE cells, both mRNA and saRNA induced innate immune responses in a RIG-I dependent manner. Since mRNA also induced innate responses, it is likely that RIG-I recognizes incoming synthetic RNAs in CHME cells. It is possible that an additional host factor such as RNA helicase modifies RNA structures of incoming synthetic RNAs delivered by LNPs and creates dsRNA, which is recognized by RIG-I and induces signals via an adaptor protein MAVS. Although KD of MAVS in 293-ISRE abrogated induction of innate immune responses to saRNAs, KD of MAVS in CHME-ISRE cells did not reduce signals although the signal is dependent on the presence of RIG-I. Currently it is unclear why KD of MAVS in CHME-ISRE cells did not impact on innate sensing of synthetic RNAs. It is possible that the KD of MAVS was insufficient, and remaining MAVS was enough to transduce signals from RIG-I in CHME-ISRE cells. Further studies including knock out of MAVS expression by the CRISPR method in CHME-ISRE cells are warranted.

We have detected robust IFN-I responses in PBMCs cultured with LNPs with saRNA. This observation is consistent with the previous report showing that synthetic RNA induces innate responses in pDCs ^8^. pDCs are well known to respond to foreign pathogens such as viruses and induce robust IFN responses ^32^. It has been shown that pDCs uptake viruses via endocytosis and endosomal RNA sensors such as TLR7/8 detect viral RNA to trigger innate immune response ^32^. It is currently unknown how pDC detects synthetic RNA formulated with LNPs. Future studies will clarify if either or both cytosolic sensors such as RIG-I and endosomal TLRs may participate in sensing. In the case of LNP-formulated mRNA, IL-1 cytokines, predominantly IL-1β, have been shown to be robustly induced in human PBMCs ^40^. In this study, we did not detect secretion of IL-1β in PBMCs stimulated with LNP-saRNA. It is possible that the differences in the platforms such as vector sequence and LNPs might have resulted in different responses. It should be noted that the published study used a relatively high dose of LNP-RNAs (5-30 µg/ml) compared to this study (50 ng/ml).

Kariko et al. reported that incorporation of 5mC in synthetic RNAs reduced innate immune responses in pDCs and DCs ^8^. Although we did not observe differences in innate immune responses in DCs to saRNA with or without 5mC, pDC produces significantly less amount of IFN-I and other cytokines in response to saRNA-5mC. We also show that 5mC-incorporated saRNA (and mRNA) had attenuated innate immune responses in THP-1 Dual cells. It is plausible that incorporation of 5mC alters secondary structures of synthetic RNAs in some cell types such as pDCs and THP-1 Dual cells, which hampers recognition of foreign RNAs by a host RNA sensor, RIG-I. Alternatively, cell-type specific expression of 5mC binding proteins may influence innate immune responses. To date, three proteins, ALYREF, YBX1 and RAD52, have been shown to recognize 5mC deposited in RNAs ^41^. Binding of these RNA modification reading proteins dictates the fate of RNA including localization and stability ^41^. It is possible that binding of such 5mC reader protein(s) to 5mC deposited on synthetic RNAs alters recognition by RIG-I potentially by blocking access of RIG-I to RNAs.

Due to the self-amplifying capacity of the saRNA vector, it is expected that the saRNA platform produces a robust and sustained expression of a target antigen. In this study, compared to mRNA, saRNA shows prolonged expression of the antigen (S RBD-TM) in both cell types tested (Fig. 1). Moreover, the injection of a small dose of saRNA vaccines in mice resulted in a robust induction of antibodies against S RBD (Fig. 6). Importantly, our phase I trial of saRNA-based SARS-CoV-2 vaccines has shown that they are safe and effectively elicit prolonged humoral immune responses against S RBD ^14^. Furthermore, this study and another clinical trial have demonstrated that an saRNA-based vaccine with 5mC induces attenuated IFN-I responses *in vitro* and reduced systemic reactogenicity when injected to human, resulting in better safety profile^15^. This saRNA-based vaccine platform, particularly with modified RNA bases, will be useful to develop vaccines with long-lasting effectiveness and reduced reactogenicity against not only SARS-CoV-2 but also other pathogens.

## Materials and Methods

### Plasmids and viruses

The saRNA plasmids encoding the nonstructural protein (nsP) 1, 2, 3 and 4 and the SARS-CoV-2 Spike RBD were constructed based on the Venezuelan Equine Encephalitis Virus (VEEV) Replicon expression vector derived from the TC-83 strain as previously reported ^42^. A lentivector expressing NanoLuc luciferase (NLuc) under the 5 tandem repeats of ISRE (pDuet 5xISRE-Nluc) was constructed by replacing ZsGreen with NLuc of Duet011 lentiviral vector expressing ZsGreen under the control of the interferon stimulated immune gene IFIT2 with a 5x repeat of the interferon-stimulated response element (ISRE) region ^43,44^. Lentiviral vectors (pLKO.1-Puro) expressing shRNA targeting MAVS, STING, RIG-I, and MDA5 were described previously ^24^. Lentiviral vectors expressing shRNA targeting MAVS, STING, RIG-I, and MDA5 and hygromycin (pLKO.1-Hygro) were generated by replacing the puromycin resistance gene of pLKO.1-Puro with the hygromycin resistance gene. A lentiviral vector expressing shRNA against MDA5 to knock down MDA5 expression in CHME-5xISRE-Nluc (see below) was purchased from Sigma (TRCN0000232948). Lentiviral particles were generated by co-transfection of HEK293T cells with lentivectors (pDuet 5xISRE-Nluc, pLKO.1-Puro, or pLKO.1-Hygro), psPAX2 (HIV packaging construct), and H-CMV-G (VSV-G expression construct)^24^. using calcium phosphate, and virus-containing cell supernatants were harvested 2 days post-transfection, cleared of cell debris by centrifugation (300 x g, 5 min), passed through 0.45 µm filters, and purified and concentrated by ultracentrifugation on a 20% sucrose cushion (24,000 rpm and 4°C for 2 hours with a SW32Ti or SW28 rotor (Beckman Coulter)). The virus pellets were resuspended in PBS, aliquoted and stored at -80 °C until use.

### Cells

HEK293T cells have been described previously ^24^. A human kidney cell line expressing luciferase under the control of the interferon-stimulated response element (ISRE) promoter (HEK293-ISRE-luc)^25^ were maintained in DMEM supplemented with 10% FBS and pen/strep with 2 µg/ml puromycin. A human microglia cell line expressing NLuc upon innate immune activation (CHME-5xISRE-Nluc) was created from a human microglial cell line (CHME3)(also known as HMC3 authenticated by ATCC ^26,27^) by transducing cells with lentiviral particles expressing 5xISRE-Nluc. Transduced cells were selected with 0.1 mg/ml hygromycin, and a single cell clone was isolated by limited dilution. Nluc expression upon innate immune activation was validated with IFN-⍺2, IFN-β, and RIG-I agonist (5’ triphosphate hairpin RNA, 3p-hpRNA, Invivogen). CHME-5xISRE-Nluc cells were maintained in DMEM supplemented with 10% FBS and pen/strep with 0.1 mg/ml hygromycin. HEK293-ISRE-luc cells or CHME-5xISRE-Nluc cells expressing shRNA against MAVS, STING, RIG-I, or MDA5 were created by transduction of cells with lentivectors expressing pLKO.1-Hygro or pLKO-Puro, respectively, followed by selection with 0.1 mg/ml hygromycin or 2 µg/ml puromycin, respectively. Cells were seeded one day prior to incubation with lipid nano particles (LNPs) containing mRNA or saRNA at 2×10^5^ cells/ml (HEK293-ISRE-luc) or 5×10^4^ cells/ml (CHME-5xISRE-Nluc) in a tissue-culture coated 96 well flat bottom plate or 12 well plate. THP-1 Dual cells (Invivogen) were maintained in RPMI1640 supplemented with 10% FBS, pen/strep, 2 mM L-glutamine, 25 mM HEPES, 100 μg/ml Normocin, 10 µg/ml of Blasticidin and 100 μg/ml of Zeocin. 1×10^5^ cells in 200 µl culture media were used for assays. Peripheral mononuclear blood cells (PBMCs) were isolated from Leukopaks (New York Biologics, Inc) using ficoll and cultured at 3×10^6^ cells/ml in RPMI1640 supplemented with 10% FBS and pen/strep (R10) after red blood cells were lysed with ammonium chloride. Monocytes (CD14+ cells) were isolated from PBMCs using anti-CD14-antibody-coated magnetic beads (Miltenyi) and cultured in RPMI1640 supplemented with 10% human AB serum (Sigma), pen/strep, and 20 ng/ml human M-CSF (Peprotech) for 5-6 days to differentiate them into monocyte-derived macrophages (MDMs). MDMs were maintained in R10 after differentiation. To generate monocyte-derived dendritic cells (MDDCs), monocytes (CD14+ cells) were cultured in R10 in the presence of human GM-CSF (10 ng/ml, Miltenyi) and IL-4 (1000 U/ml, BD) for 5 days as previously described ^24^. Plasmacytoid dendritic cells (pDCs) were isolated from PBMCs using human Plasmacytoid Dendritic Cell Isolation Kit II (Miltenyi) or human CD304 (BDCA-4/Neuropilin-1) MicroBead Kit (Miltenyi).

### saRNA synthesis and lipid nano particle (LNP) formulation

Plasmids with membrane-anchored SARS-CoV-2 S-RBD (Gamma) ^10^ with or without PADRE, or GFP placed downstream from a T7 promoter were linearized by digestion with the BspQ1 restriction enzymes at 50 °C for 3 h. The linearized plasmid was then purified using the Wizard Plus SV Miniprep DNA Purification System (Promega), and saRNA was transcribed in vitro using HiScribe T7 High Yield RNA Synthesis Kit (NEB). After DNase treatment, the saRNA was purified with RNeasy midi kit (Qiagen), and subsequently modified by the addition of a 7-methylguanosine cap with the Vaccinia Capping System (New England Biolabs [NEB]) using the NEB Capping protocol (NEB, M20280). After purification of the capped saRNA using the Monarch kit (NEB), LNP-saRNA was formulated with GenVoy-ILM (Precision Nano Systems) with the NanoAssemblr Ignite (Precision Nano Systems).

### LNP culture

LNPs were diluted to 25 ng/µl with 4% sucrose in PBS, aliquoted in a small volume and stored at -80°C until use. LNPs were serially diluted (5-fold) with culture media starting with 50 or 500 ng/ml then cultured with cells. As a positive control, cells were transfected with a RIG-I agonist (5’ triphosphate hairpin RNA, 3p-hpRNA, Invivogen). Cells were analyzed for antigen expression or luciferase/NLuc expression, and culture supernatants were analyzed for cytokine expression as follows.

### Luciferase/NLuc assay

Cells in a 96 well plate were lysed with Glo-Lysis buffer (Promega, 50 µl) and 25 µl cell lysates were mixed with 25 µl Bright-Glo or Nano-Glo (both Promega), and luminescence was measured by a luminometer. Culture supernatants from THP-1 Dual cells (20 µl) were mixed with QUANTI-Luc 4 Lucia/Gaussia reagent (Invivogen), and luminescence was measured by a luminometer. Values were normalized to those in no-treatment samples (media without LNPs).

### Flow cytometry (FCM)

For the detection and quantitation of cell surface expression of SARS-CoV-2 S protein RBD (S-RBD), cells were detached with enzyme-free cell dissociation buffer (Cellstripper, Corning) and stained with live/dead dye (Zombi NIR, BioLegend, 1:250 in PBS) for 20 min at RT. After quenching dye with 2% NCS in PBS (washing buffer), cells were stained with human anti-SARS-CoV-2 S antibody (M128, Acro Biosystems, 1:200) for 30 min at 4°C, followed by washing with washing buffer twice, and stained with a secondary anti-human IgG-PE (1.5 µl per sample) for another 30 min at 4°C. After washing, cells were fixed with 4% PFA. PBMCs were harvested and stained with live/dead dye (Zombi NIR, 1:250 in PBS) for 20 min at RT. After quenching dye with 2% NCS in PBS (washing buffer), cells were incubated with Fc blocker (2.5 µl in 20 µl, BD) for 10 min at RT then stained for the following cell surface markers for 30 min at 4°C: CD3 (T cell marker, mouse anti-CD3-BV510, BioLegend, 1:50), CD14 (monocyte marker, mouse anti-CD14-BV421, BioLegend, 1:50), CD19 (B cell marker, mouse-anti-CD19-PE-Cy5, 1:25), CD169 (myeloid cell activation marker, mouse anti-CD169-Alexa647, BioLegend, 1:100), HLA-DR (as an activation marker, mouse anti-HLA-DR-PE, BD, 1:25) in the presence of True-stain (BioLegend, 1:20) to prevent non-specific binding to certain fluorophores such as PE-Cy5. After washing, cells were fixed with 4% PFA. MDMs were detached from a plate using an enzyme-free cell dissociation buffer (Corning) and stained with a live/dead dye and anti-CD169-Alexa647 antibodies as described above. All samples were analyzed with a BD LSRII flow cytometer, and data were analyzed with FlowJo software (FlowJo).

### Multiplex cytokine assay

Culture supernatants from cells with LNPs were harvested on Day 1, 2, and 3, and stored in -80°C until use. Cytokine release in the supernatants was measured with an Inflammation 20-Plex Human ProcartaPlex™ Panel (Invitrogen) and was analyzed with the DropArray (Curiox Biosystems) and the MAGPIX System (Luminex) or with LEGENDPlex Human Anti-Virus Response Panel (13-plex, BioLegend) and was analyzed with LSRII (BD) or Aurora (Cytek). Data was analyzed with FlowJo software (FlowJo) or the LEGENDplex Data Analysis Software Suite (BioLegend).

### mRNA Quantification

Total mRNA was isolated from 5×10^5^ cells using a kit (RNeasy kit, QIAGEN) and reverse-transcribed using oligo(dT)20 primer (SuperscriptIII, Invitrogen). Target mRNA was quantified using Maxima SYBR Green (Thermo Scientific) and normalized to GAPDH mRNA by the 2-ΔΔCT method ^45^.

### Immunoblot Analysis

To validate knockdown of host proteins (MAVS, STING, RIG-I and MDA5), cell lysates containing 20-30 µg total protein were separated by SDS-PAGE, transferred to nitrocellulose membranes and the membranes were probed with the following antibodies: a rabbit anti-MAVS polyclonal antibody (Thermo Fisher), a rabbit anti-STING polyclonal antibody (Cell Signaling), a mouse anti-RIG-I antibody (Enzo Life Sciences) or a rabbit anti-MDA5 antibody (Proteintech). Then, the membranes were stained with secondary antibodies, a goat anti-mouse-IgG-DyLight 680 (Pierce) or a goat anti-rabbit-IgG-DyLight 800 (Pierce). As loading controls, actin was probed using a rabbit anti-actin antibody (SIGMA) or a mouse anti-actin antibody (Thermo Fisher). Membranes were scanned with an Odessy scanner (Li-Cor).

### Enzyme-Linked Immunosorbent Assay (ELISA)

ELISA was used to evaluate anti-SARS-CoV-2 antibodies in immunized mice. Nunc-Immuno MicroWell 96-well plates (Thermo Fisher Scientific) were coated with recombinant SARS-CoV-2 (2019-nCoV) Gamma RBD (Sino Biological) at 0.1 µg/mL in PBS overnight at 4°C. Coated plates were washed 3 times with Tris-buffered saline with 0.05% Tween-20 (TBS-T) before blocking with blocking buffer (5% non-fat milk in TBS-T) for 1 hour at ambient temperature. After blocking, serum samples were plated at a 1:50 initial dilution, followed by 3-fold serial dilution in blocking buffer. The plates were washed again 3 times before addition of 100 µL of the appropriate HRP-conjugated secondary antibody. After one hour-incubation at room temperature, the plates were washed, developed using 3, 3’, 5, 5’-tetramethylbenzidine (TMB) Microwell Peroxidase Substrate (Seracare) for 20 minutes, and stopped by 2N sulfuric acid in water (Avantor). The absorbance values at 450nm (OD_450_) were read in BioTek Synergy HTX Multi-Mode reader (Agilent). Endpoint tiers were defined as the log_10_ dilution resulted in an OD_450_ = 0.5 using 4PL regression and GraphPad Prism software. Serum titers with undetectable binding were assigned the value one-half lower limit of detection (LLOD), while those over OD_450_ = 0.5 were assigned the value upper limit of detection (ULOD). IP-10 in tissue culture supernatants was quantified with a Human IP-10 ELISA Set (BD).

### Animal Study

The immunogenicity study in mice was conducted at Noble Life Sciences Inc, (Sykesville, MD). Female 6- to 8-week-old Balb/c mice (n = 6 per group) were injected intramuscularly with 10 µg of the saRNA expressing S RBD-TM formulated in lipid nano particle in 50 µL of Dulbecco’s phosphate buffered saline (DPBS) containing 4% sucrose at weeks 0 and 4. An equivalent volume of DPBS was injected for the control group. Mouse sera were collected at day 0, 14, 21, 29, 42, and 56.

## Acknowledgement

We would like to acknowledge the technical support from the Boston University Medical Campus Analytical Core and the Flow Cytometry Core. This work was supported by National Institute of Health (NIH) grants R01AG060890 (S.G.), R01DA051889 (S.G.), R01DA055488 (S.G.), P30AI042853 (S.G. and H.A.), T32AI007309 (J.F.), F31AI172625 (J.F.), and R21NS126092 (H.A.).

## Author contributions

W.A. and H.A. conceived, designed, and coordinated the study. M.K., A.L.M., L.R., E.P., A.A.Q, J.F., and H.A. performed the experiments. K.M., J.F.S., W.A., S.G. and H.A. contributed to the writing and revision of the manuscript. All authors read and approved the manuscript.

## Conflict of interest

M.K., A.L.M., L.R., E.P., and K.M. are employees of VLP Therapeutics, Inc.; J.F.S. is an employee and holds stocks in VLP Therapeutics, Inc.; W.A. is a board member, an employee and holds stocks in VLP Therapeutics, Inc. and is a management board member of VLP Therapeutics Japan, Inc.; W.A. and J.F.S are inventors on related vaccine patent. The remaining authors declare no competing interest.

## Figure Legends

**Supple. Fig. 1.**
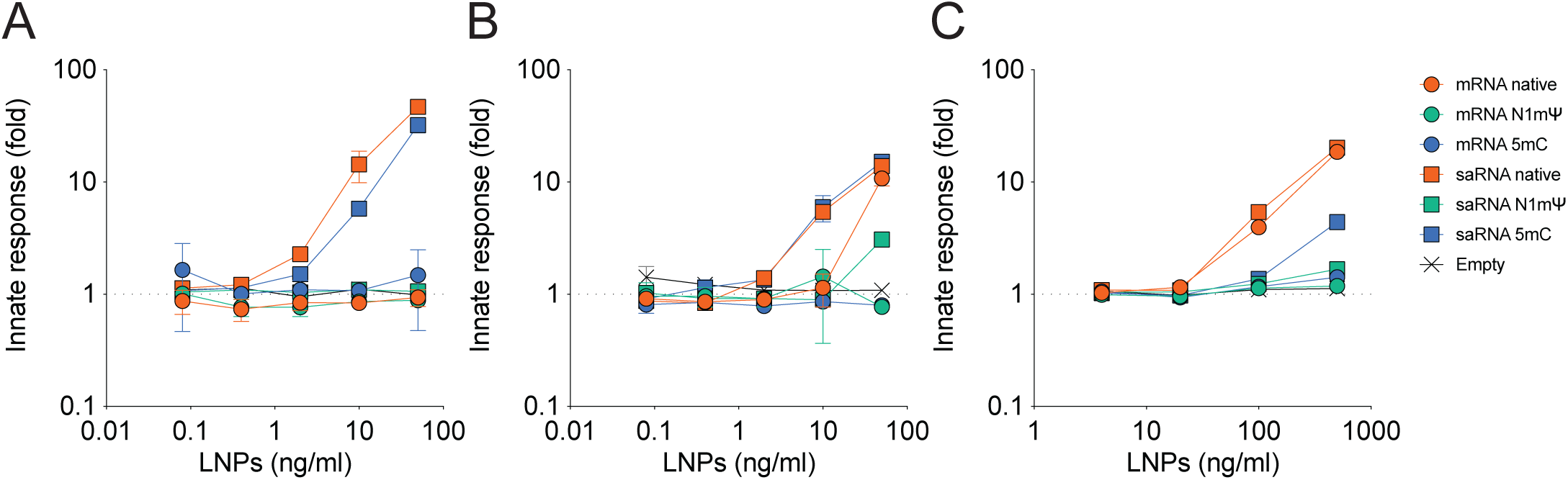
LNP-formulated mRNA and saRNA induce innate immune activation in a dose dependent manner. (A) 293-ISRE cells, (B) CHME-ISRE cells, or (C) THP-1 Dual cells were incubated with increasing amounts of with LNPs containing mRNA or saRNA with native bases, N1mΨ, or 5mC, or empty LNPs and luciferase or Nluc activity was quantified 24-48 hours post stimulation. Values were normalized to those from media-only control. Representative graphs are shown (n=4).

**Supple. Fig. 2.**
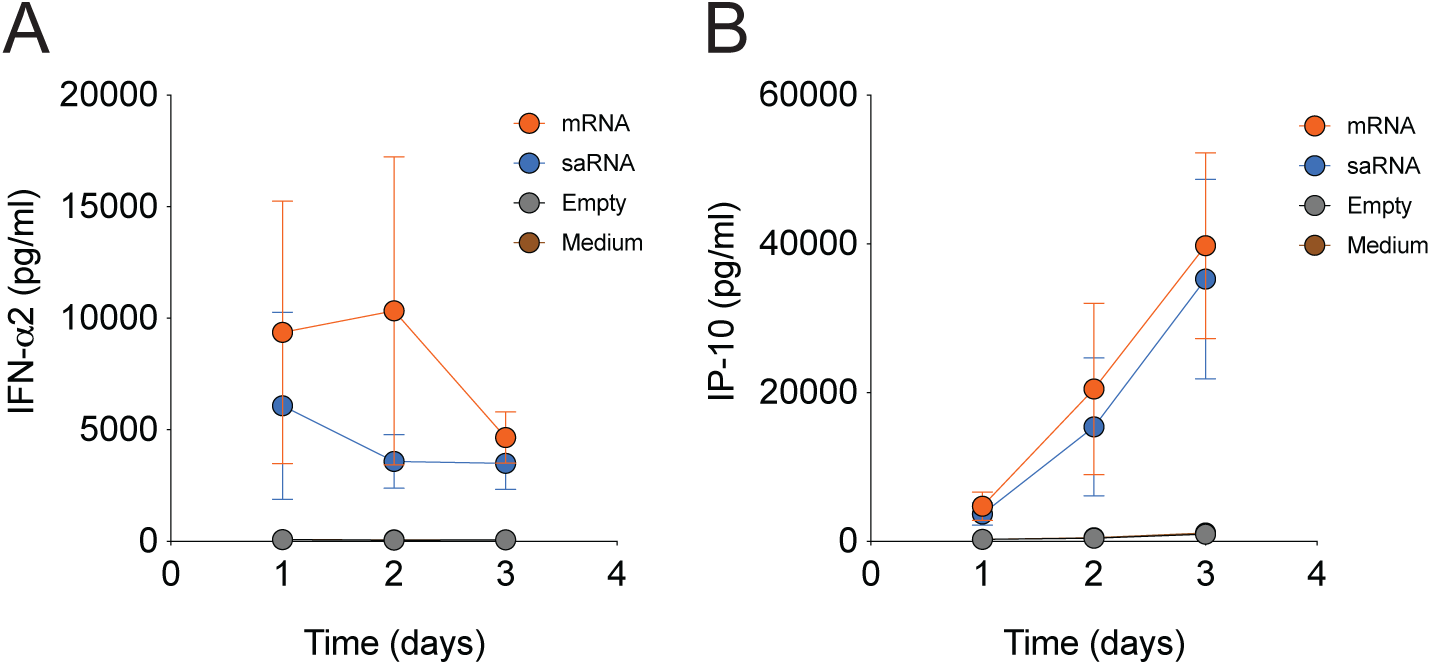
Kinetics of IFN-⍺2 and IP-10 production in PBMCs. PBMCs from healthy donors were incubated with mRNA-native (mRNA), saRNA-native (saRNA), or empty LNPs (Empty) at 50 ng/ml, culture supernatants were harvested 1-3 days post initiation of culture, and (A) IFN-⍺2 and (B) IP-10 production in the sups were quantitated with multiplex cytokine assay (Luminex). The graph shows the means +/- SEM (n=6).

**Supple. Fig. 3.**
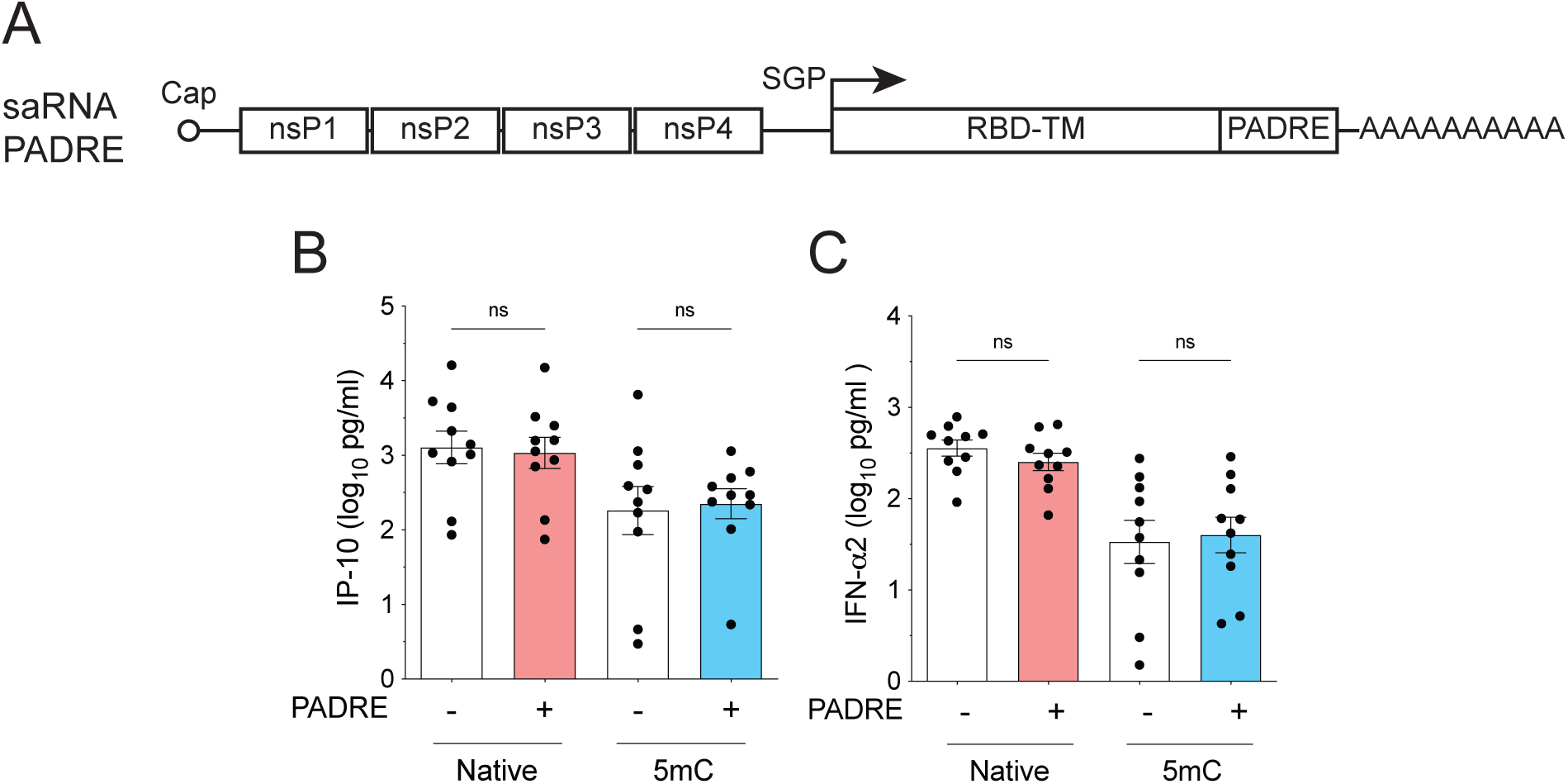
The saRNA-PADRE vector induces same levels of innate immune responses in PBMCs as the saRNA vector. (A) Schematics of the saRNA-PADRE vector expressing membrane-anchored SARS-CoV-2 S-RBD (RBD-TM). SGP: subgenomic promoter. (B and C) PBMCs were cultured with LNPs containing saRNA (-) or saRNA-PADRE (+) with native or 5mC at 50 ng/ml for 1 day, and (B) IP-10 and (C) IFN-⍺2 released into the culture were analyzed by multiplex cytokine assay (LegendPlex). One-way ANOVA, ns: not significant.

**Supple. Fig. 4.**
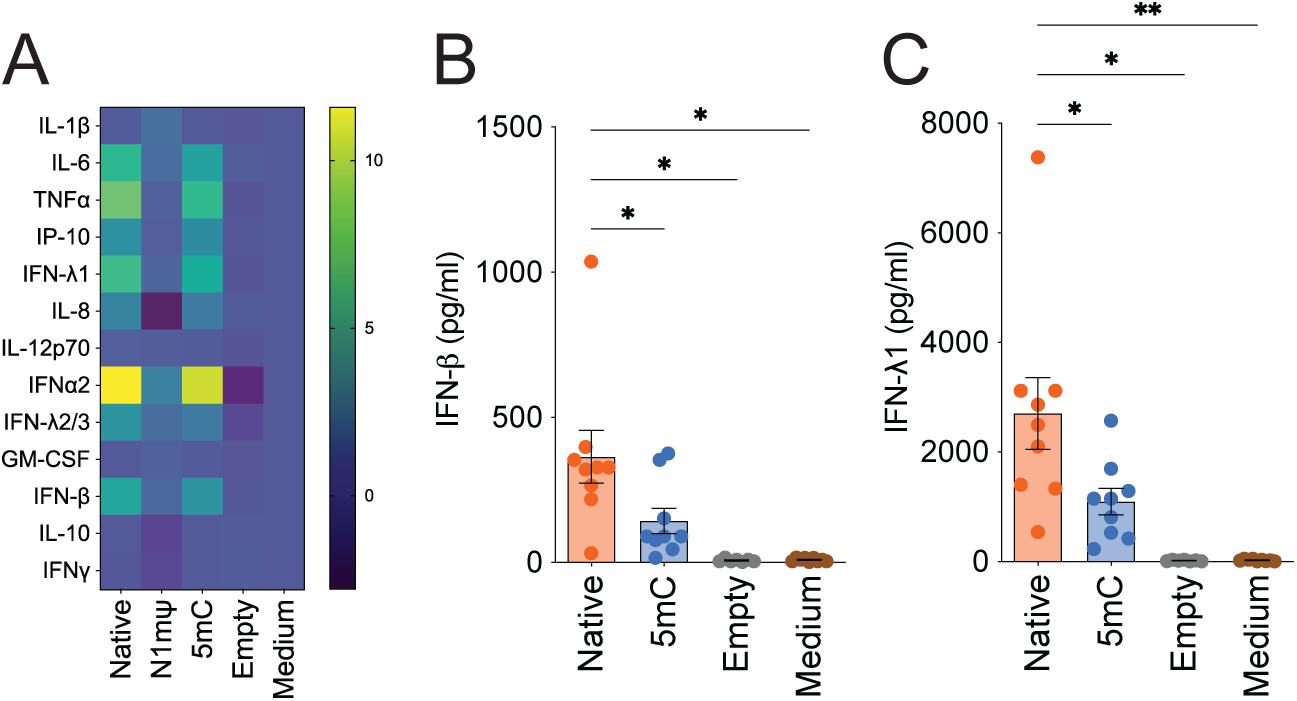
pDCs induce a variety of proinflammatory responses to saRNA, which is attenuated by 5mC incorporation. (A) pDCs isolated from healthy donor PBMCs were incubated with LNPs containing saRNA-PADRE (native, N1mΨ, 5mC), or empty LNPs at 50 ng/ml for 1 day, and cytokines released into the sups were quantitated with LegendPlex. Values were normalized to those of medium only control, and fold change in log_10_ is shown. (B) pDCs were incubated with LNPs containing saRNA-PADRE (native or 5mC), or empty LNPs at 50 ng/ml for 1 day, and the production of (B) IFN-β or (C) IFN-λ1 was measured with LegendPlex. One-way ANOVA, *: *p*<0.05, **: *p*<0.01.

**Supple. Fig. 5.**
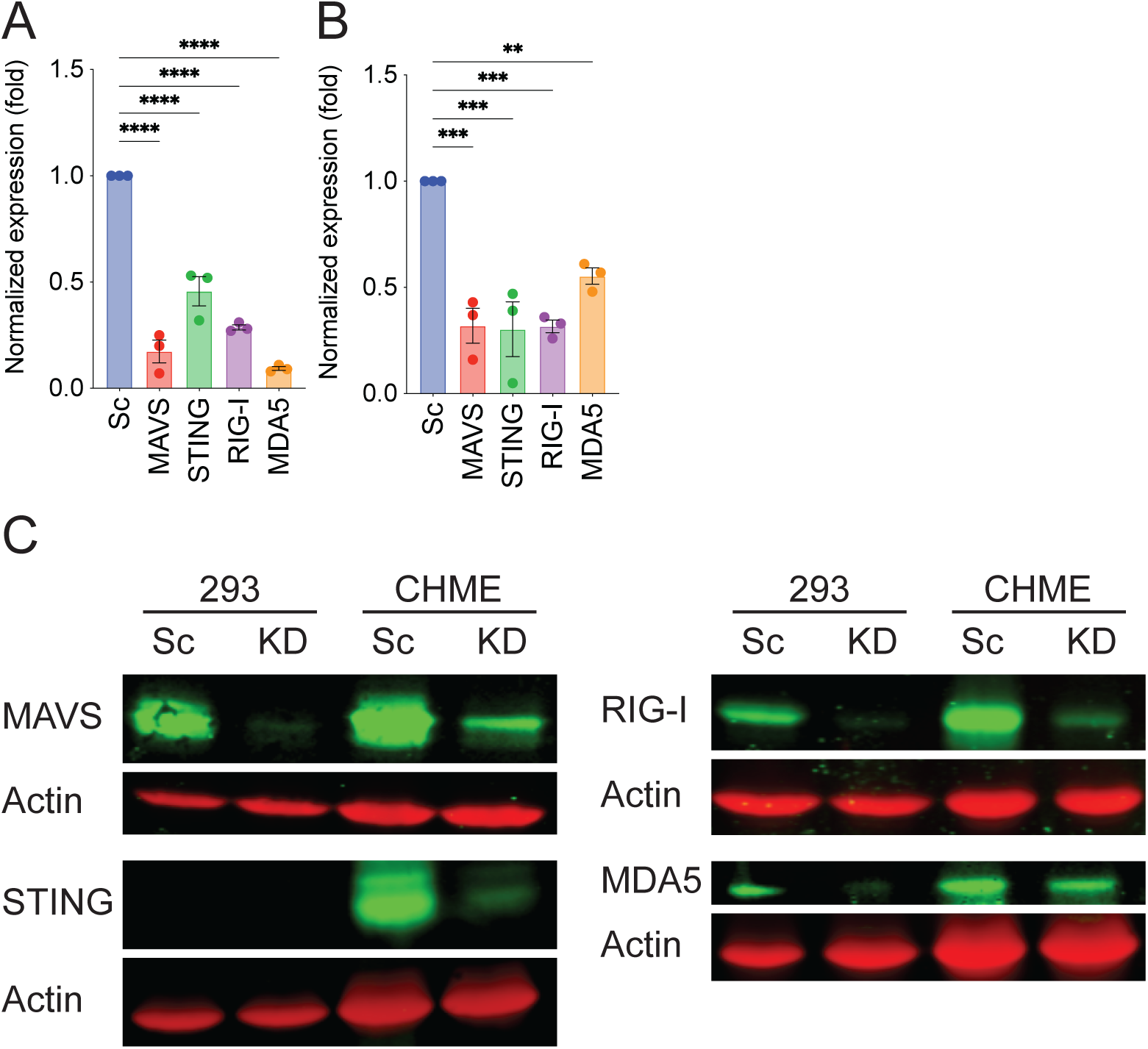
KD of RIG-I and MAVS abrogated innate immune activation against synthetic RNAs. 293-ISRE cells and CHME-ISRE cells were transduced with lentivectors expressing shRNA targeting the indicated genes and selected for successful transduction using puromycin or hygromycin. KD efficiency in (A) 293-ISRE cells and (B) CHME-ISRE cells was quantified by qRT-PCR. The gene expression was normalized to that in cells with scrambled shRNA. Each dot represents an individual experiment, and the graph shows the means +/- SEM. (C) Western blotting analysis of protein expression in 293-ISRE cells (293) or CHME-ISRE cells (CHME). Sc: Scrambled shRNA expressing cells, KD: KD cells. Actin was used as an internal control. One-way ANOVA, **: *p*<0.01, ***: *p*<0.001, ****: *p*<0.0001.

